# Development of zebrafish (*Danio rerio*) as an *in vivo* model for *Borrelia burgdorferi* infection

**DOI:** 10.1101/2022.06.30.498276

**Authors:** Erica Misner, Min Zhang, Eva Sapi

## Abstract

*Borrelia burgdorferi* is the spirochetal bacterium that causes Lyme disease. Despite the fact that antimicrobial sensitivity of *B. burgdorferi* has been widely studied, there is still a need to develop an affordable, practical, high-throughput *in vivo* model which can be used to find effective antibiotic therapies, especially for the recently discovered persister and biofilm forms. Recent studies showed that Zebrafish (*Danio rerio*) could offer a novel, high-throughput, affordable model in antibiotic therapies for various infections agents can be studied. Therefore, in this study, we developed a straightforward, standardized infection procedure and tested it for zebrafish survival rate, morphological or behavioral changes post-infection as well as providing evidence that *B. burgdorferi* persists in zebrafish using polymerase chain reaction (PCR) and immunohistochemical (IHC) techniques. Morphological and physiological examination showed a significant size difference between control and infected zebrafish at 2 wpi, and infected zebrafish exhibited slower heart rates through 72 hpi. Furthermore, our results showed *B. burgdorferi* DNA can be detected and active replication of the *B. burgdorferi* 16S rRNA gene can be confirmed through 10 days post-infection via PCR and Reverse Transcription PCR respectively. Fluorescent microscopy and immunohistochemical staining revealed spirochetes present in the eyes, gills, heart, liver, tail, and hindbrain tissues though 72 hpi as well as the stomach and digestive tract at 2 weeks post-infection, respectively. These findings demonstrate that zebrafish could serve as a promising animal model to study the mechanism of *B. burgdorferi* infection as well as *in vivo* antibiotic sensitivity.

## Introduction

*Borrelia burgdorferi*, the causative agent of Lyme disease, is estimated to infect and cause Lyme disease in 476,000 people every year, making it the number one vector-borne illness in the United States [1]. Lyme disease has a broad range of symptoms, including arthritis, Lyme carditis, lymphocytoma, meningitis and other neurological indispositions, and can affect everyone differently [2,3]. The most well-characterized morphological form of *Borrelia* is the spirochete, though it can change its morphology based on its environment, available nutrients, and stress factors [2,4,5,6]. Spirochetes are the actively metabolizing, motile form – utilizing flagella to evade host immune response and disseminate to various tissues of the body. *B. burgdorferi* spirochetes have been found in the kidneys, liver, heart, and spinal cord of human subjects [5,7,8], in the brain of non-human primates [8,9], and in the heart, brain, joints, and muscle tissue of mice [7,9,10]. Alternative forms such as *B. burgdorferi* round bodies form as a result of unfavorable pH, low nutrients, high oxygen, presence of antibiotics, and other factors [4,5,11,12] and it was shown that they have high resistance to several antibiotics, such as doxycycline [13,14], ceftriaxone [9], and tigecycline [15].

However, there is another form of *B. burgdorferi* found and characterized, called biofilms, which are spirochetal aggregates protected by a complex extracellular polymeric layer [6,14]. Biofilms, such as those produced by *Staphylococcus aureus* and *Pseudomonas aeruginosa* [16] are known to be extremely resistant to antibiotics and are strong contributors to chronic diseases [17]. Studies which have addressed antibiotic resistance of *B. burgdorferi* biofilm form showed similar high antibiotic resistance [14] and only a powerful combination of dapsone/doxycycline/rifampin/azithromycin had some effect on *B. burgdorferi* biofilms, at least *in vitro* [18]. There is an obvious need to have an animal model for *in vivo* studies which can affordably test all these antibiotic combinations for different species of *B. burgdorferi* sensu lato.

Animal models such as mice and rhesus macaques have been instrumental in providing information about Lyme disease and are more representative of the human body than *in vitro* studies. Unfortunately, testing antibiotics in these models is time consuming and very expensive [9,13,19]. The arsenal of antibiotic treatments tested *in vivo* so far is by no means exhaustive and calls for further research to discover more effective combinations against *B. burgdorferi*. Therefore, we have proposed the zebrafish (*Danio rerio*) as a new animal model to study *B. burgdorferi* infection and antibiotic testing [20]. Antibiotic susceptibility assays can be performed easily by immersing zebrafish directly in antibiotic suspensions and observing the effects, for example, on *Mycobacterium abscessus* or *P. aeruginosa* infections [21,22]. Zebrafish are affordable and effective models due to their small size, simple housing and maintenance, and high throughput capabilities thanks to their prolific breeding capacity [23]. As embryos, their transparency allows for pigmentation to be delayed enabling visualization of organ growth and real-time bacterial dissemination [24,25]. Having over 70% genetic similarity to humans [26], zebrafish react to bacterial infections with adaptive and innate immune responses parallel to mammals, including IL-1, IL-6, TNFa, TLR5, and other inflammatory cytokines, macrophages/neutrophils, and natural killer cells [20,23,27,28].

To validate zebrafish as a suitable Lyme disease model, we subjected 5-day-old zebrafish larvae to suspensions of media with or without *B. burgdorferi* spirochetes for 8 hours. The infected zebrafish were observed and evaluated for morphological and physiological changes. The success of infection and the persistence *B. burgdorferi* in zebrafish were evaluated by fluorescence immunohistochemical staining, PCR and Reverse Transcription PCR for 2 weeks post-infection.

## Materials and methods

### Ethics statement and zebrafish breeding

All experiments were conducted according to the protocol approved by the Institutional Animal Care and Use Committee (IACUC Protocol No. 19-02), University of New Haven (New Haven, CT). All handlers conducting research on the zebrafish underwent IACUC and CITI program training. Adult zebrafish used in this study (wildtype TUAB) were housed in tanks receiving continuously filtered fresh water from the Iwaki Zebrafish Research System at the University of New Haven. Lights were on a strict 14-hour daylight cycle. Up to four male and four female zebrafish were kept as pairs in breeding tanks overnight. In the morning, dividers were removed, and pairs were allowed to breed for up to two hours. Embryos were then incubated at 28°C in Petri dishes containing water from the Iwaki System supplied with 0.0002% methylene blue (Sigma-Aldrich, St. Louis, MO, USA) to prevent fungal contamination. Embryo water was replaced daily, and larvae were reared until 5-days post-fertilization (dpf) where they could then be used for immersion infection. Fig. 1 shows a timeline for the use, infection, and processing of zebrafish at different stages, including parent adult through offspring larvae. No zebrafish larvae (uninfected or infected) were used beyond the allotted experimental period; at the end of each experiment (between 24 hours post-infection and 2 weeks post-infection), larvae were euthanized by tricaine overdose then immediately processed by genomic extraction or fixation as described below.

**Figure 1.**
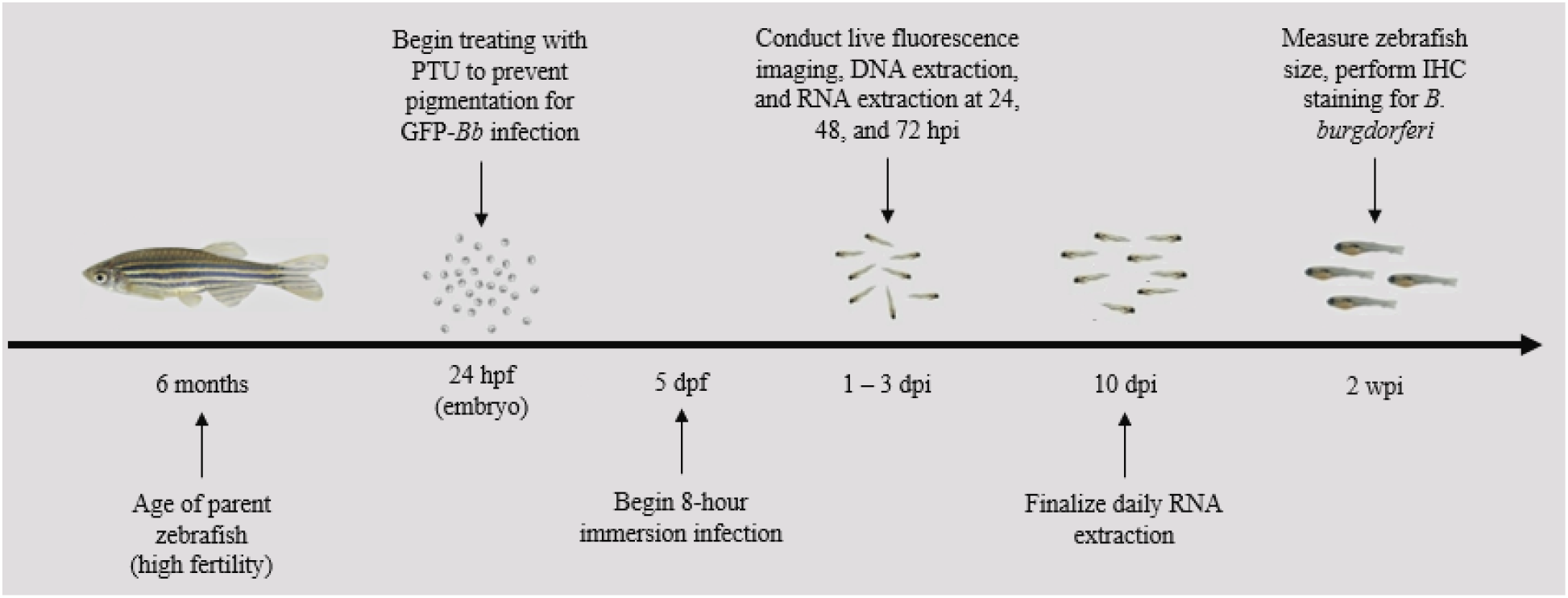
Timeline of use for zebrafish including adult breeders and offspring larvae.

### *B. burgdorferi* culture

Two strains of *B. burgdorferi* sensu stricto were utilized for immersion infection. To visualize *B. burgdorferi in vivo*, low passage isolates (passage number < 6) of a genetically engineered green fluorescent B31 strain (GFP-B31) from Dr. G Chaconas (University of Calgary; [29] were cultured using sterile 15-mL glass tubes with modified Barbour-Stoenner-Kelly (BSK-H) media (Sigma-Aldrich, B3528) supplemented with 6% rabbit serum (Pel-Freez, 31125-5) and 10 ug/mL gentamicin (Sigma-Aldrich, 1405-41-0) to maintain GFP-tagged *B. burgdorferi*. For increased pathogenicity, a more virulent strain, *B. burgdorferi* N40, was cultured under the same conditions, omitting gentamicin. Both strains were incubated at 33°C with 3% CO_2_ until reaching log-phase.

### Immersion infection

Zebrafish larvae which were free from the chorion at 5 dpf were used for immersion infection. Larvae were fed with GEMMA size 75 (Skretting, Tooele, UT, USA) zebrafish food 1 hour prior to infection. For each infection group, 20 larvae were moved to a Petri dish and washed twice with 20 ug/mL gentamicin. Under the hood, gentamicin was removed and replaced with 6 mL of autoclaved fish water from the Iwaki System for both control and experimental dishes. Into the control dish, 6 mL of sterile BSK-H media (Sigma-Aldrich) were added. For the experimental suspension, a log-phase culture of *B. burgdorferi* sensu stricto strain N40 was checked for motility and confluence, counted, then pelleted at 3200 rpm for 15 minutes at room temperature. The pellet was resuspended in 6 mL of media, then added to the experimental dish to achieve a final concentration of 1×10^7^ spirochetes/mL. Both control and experimental fish were incubated at 28°C for 8 hours. Following the 8-hour infection, darkfield microscopy was used to verify that the control suspension contained no *B. burgdorferi*, while experimental suspension contained motile spirochetes. Under the hood, all media was removed from each dish and fish were washed twice with media containing 20 µg/mL gentamicin, then once with sterile fish water. Zebrafish were returned to the incubator at 28°C until 24 hpi. Fish were fed twice daily and received daily water changes.

To visualize fluorescent spirochetes in the fish between 24 and 72 hpi, an alternative method of infection was used. Starting at 24 hpi, zebrafish embryos were treated with 180 µM of 1-phenyl 2-thiourea (PTU) (Sigma-Aldrich, 103-65-5), to prevent pigmentation [25]. The PTU-embryo water suspension was changed daily through 72 hpi. Immersion setup starting at 5 dpf remained the same as *B. burgdorferi* N40 infection; however, gentamicin was diluted with autoclaved zebrafish water into 20 ug/mL suspension to prevent loss of the GFP tag throughout infection [30]. In each dish, 6 mL of the 20 µg/mL gentamicin was mixed with 6 mL of BSK-H (with or without GFP-Borrelia) to obtain a final suspension of 10 µg/mL gentamicin containing 1×10^7^ GFP-Borrelia spirochetes. All experiments involving GFP-Borrelia were done in the dark to prevent loss of spirochete fluorescence.

### Real-time monitoring and fluorescence microscopy

Beginning at start of infection (*t* = 0), end of infection (*t* = 8 hours), and twice daily through 72 hpi, overall health and appearance of zebrafish were observed, including any changes to behavior such as lethargy or flashing against the bottom of the dish were noted. Zebrafish which displayed discoloration or were unresponsive to stimuli were removed from the infection group and euthanized by tricaine methanesulfonate overdose (300 mg/L) for 20 minutes between 0- and 12-hours post-onset of endpoint criteria. Zebrafish mortality occurring prior to the end of the observation period was due to natural causes by bacterial burden.

Survival rates were recorded at 24, 48, and 72 hpi. Size and morphological differences were evaluated between uninfected and infected zebrafish between 24 and 72 hpi and at 2 wpi. To measure size, zebrafish anesthetized with 200 µg/mL Tricaine methanesulfonate (MS222) (Sigma-Aldrich, 886-86-2) were imaged under a ZEISS Stemi 508 stereo microscope at 3.5x (Carl Zeiss Microscopy, LLC. NY, USA) and measured across a computer monitor using a ruler for more precise measurements. Additionally, heart rates were quantified between infection groups. Here, each day through 72 hpi, 5 out of 20 zebrafish were randomly selected and moved to Petri dishes containing sterile water and left to acclimate under a stereo microscope. The heart rate of each fish was measured for 60 seconds, following which all fish were returned to the incubator.

To visualize the presence of GFP-Borrelia spirochetes at 24, 48, and 72 hpi, 20 uninfected and 20 infected zebrafish were anesthetized for 15 minutes with 200 µg/mL Tricaine to reduce stress. Fish were imaged in the dark using a ZOE Fluorescent Cell Imager, then returned to fresh autoclaved fish water. For visualization of spirochetes at 2 wpi, zebrafish were euthanized, fixed in 4% paraformaldehyde (PFA) at 4°C overnight, followed by 20% EDTA decalcification at room temperature for three days. Fish were rinsed with two 20-minute nuclease-free water washes, then placed in plastic histology cassettes for soaks. Fish tissues were subjected to a dehydration series of increasing ethanol: 30%, 50%, 70%, and twice in 100% ethanol for 20 minutes each. Cassettes underwent a series of 20-minute infiltration steps: 1:1 ethanol-xylene, 1:3 ethanol-xylene, then 100% xylene. Cassettes were left in a glass dish (set in a 60°C water bath) containing 1:1 xylene-paraffin for 1 h, then again in 100% paraffin overnight. Finally, tissues were embedded horizontally in molten paraffin. Embedded zebrafish blocks (both infected and uninfected) were sectioned at 4 μm, deparaffinized, then stained with a *Borrelia* polyclonal antibody diluted to 1:50 in 1x PBS (ThermoFisher, Waltham, MA, USA; FITC-PA1-73005) according to a previously established protocol [31]. Negative control slides omitted the primary *Borrelia* antibody. Slides were visualized at 400x with a Nikon Eclipse 80i fluorescent microscope (Nikon Metrology, Inc., Tring, UK).

### Wholemount Immunohistochemistry

To visualize non-GFP *B. burgdorferi* inside the zebrafish, larvae were processed by wholemount immunohistochemistry (IHC) at 24, 48, and 72 hpi. From each group (uninfected and infected), 4 zebrafish were selected, euthanized by tricaine overdose, then immediately fixed in 500 µL of 4% PFA at 4°C overnight. Whole-mount immunohistochemistry was performed using a reference protocol [32] with the following modifications. After the blocking with 10% BSA/PBST at 4°C overnight, the zebrafish were incubated with 200 µL of 1:100 primary *B. burgdorferi* monoclonal antibody (ThermoFisher, F21l) diluted in 1% normal goat serum (NGS) at 4°C for 24 hours. The zebrafish were then washed 4x with 500 µL PBST for 30 minutes. The next day, the primary antibody was removed, and the zebrafish were washed 4x with 500 µL PBST for 30 minutes. For each zebrafish, 200 μL of 1:100 secondary antibody with a fluorescent tag (goat anti-mouse IgG1, DyLight 594 conjugated) diluted in 1% NGS were added, then the zebrafish were incubated at room temperature for 4 hours in the dark. The zebrafish were washed 5x with PBST for 5 minutes Finally, the zebrafish were immersed in 500 µL of 25%, 50%, 75%, 100% glycerol, for 20 minutes for each concentration. The whole-mount IHC zebrafish were observed using a Leica DM2500 fluorescent microscope (Leica Microsystems, Inc., Buffalo Grove, IL, USA) at 100x magnification. Uninfected zebrafish served as negative control.

### PCR analysis

#### DNA detection via nested PCR

To assess the ability of *B. burgdorferi* to infect zebrafish, and to ensure that they were still harboring spirochetes through 72 hpi, DNA extraction was conducted at 24, 48, and 72 hpi. At each of the timepoints, 8 zebrafish were transferred to sterile 1.5-mL microfuge tubes using transfer pipettes, then euthanized by Tricaine overdose and washed twice with 1x PBS. A DNeasy Genomic DNA Extraction Kit (QIAGEN, Hilden, Germany; #69504) was used to extract DNA according to the manufacturer’s instructions.

For DNA amplification, a nested PCR protocol for *B. burgdorferi* detection was followed [33], which used *PyrG* (CTP synthase) primers and HotStarTaq PCR reagents (QIAGEN, #203203). DNA samples were quantified then diluted to 20 ng/µL so that 5 µL could be used in each 25-μL PCR reaction. Each sample contained the following: 1x HotStarTaq buffer, 1.5 mM MgCl_2_, 0.2 mM dNTP mix, CTP synthase primers at 0.4 µM final concentration each (Forward: 5’ ATTGCAAGTTCTGAGAATA; Reverse: 5’ CAAACATTACGAGCAAATTC), 2.5 units of HotStarTaq polymerase, 5% DMSO, 100 ng of template DNA, and PCR-grade water up to 25 µL. Samples were run in a T100 thermal cycler under the following protocol (Bio-Rad, Hercules, CA, USA): initial denaturation at 94°C for 15 minutes; 30 cycles of 94°C/30 seconds, 55°C/30 seconds, 72°C/30 seconds; and final extension at 72°C for 5 minutes. Round 1 PCR products were then diluted 1:100 in molecular-grade water prior to use in round 2.

Round 2 nested PCR samples were prepared by mixing each of the following: 1x HotStarTaq buffer, 2.5 mM MgCl_2_, 0.2 mM dNTP mix, nested primers each at 0.4 µM (Forward: 5’ GATATGGAAAATATTTTATTTATTG; Reverse: 5’ AAACCAAGACAAATTCCAAG), 1.5 units of HotStarTaq polymerase, 1% formamide, 5 µL of diluted template DNA, and PCR-grade water up to 50 µL. Samples were run in a Bio-Rad T100 thermal cycler under the following protocol: initial denaturation at 94°C for 15 minutes; 35 cycles of 94°C/30 seconds, 50°C/30 seconds, 72°C/30 seconds; and final extension at 72°C for 5 minutes. PCR products were analyzed by standard agarose gel electrophoresis in a 1% gel referenced against a Hi-Lo DNA ladder (Bionexus, Inc., Oakland, CA, USA; BN2050). An infected zebrafish sample which was positive for *B. burgdorferi* was purified using a GeneJET PCR Purification Kit (Thermo Scientific, #K0701), then sent for Eurofins sequencing and analyzed using the NCBI BLAST program (https://blast.ncbi.nlm.nih.gov/). Negative controls included a no template-control which received nuclease-free water instead of DNA template as well as uninfected zebrafish samples.

### RNA detection via RT-RT PCR

To verify active *B. burgdorferi* transcription inside infected zebrafish larvae from 1 through 10 dpi (and no transcription in uninfected zebrafish), RNA extraction was performed followed by reverse-transcriptase PCR. For each sample, RNA was extracted from either 3 uninfected or 3 *B. burgdorferi*-infected zebrafish following manufacturer’s instructions for the RNeasy mini kit (QIAGEN, #74104).

All RNA samples were quantified and followed by cDNA synthesis using a First-Strand cDNA Synthesis Kit for Real-Time PCR (USB-Affymetrix, Santa Clara, CA, USA; #75780). Samples were prepared according to manufacturer’s instructions for a 20-µL reaction, each containing 2 µL of primer mix, 2 µL of 10X RT buffer, 1 µL of 10 mM dNTPs, 1 µL of RNase inhibitor, 1 µL of M-MLV RT, total RNA up to 1 µg, and RT-PCR grade water up to 20 µL. The cDNA synthesis reactions were run on a Bio-Rad T100 thermal cycler under the following protocol: 44°C for 60 minutes; then 92°C for 10 minutes. Next, each cDNA sample was amplified for *B. burgdorferi* 16S rRNA by preparing samples in a 96-well plate using 5 µL of 2X SYBR Green Master Mix (Bio-Rad #1725121), 1 µL of each primer (Forward: 5’ GTGGCGAACGGGTGAGTAAC; Reverse: 5’ CCGTCAGCTTTCGCCATTGC), 100 ng of cDNA, and RT-PCR grade water up to 10 µL. A BioRad CFX96 RealTime System was used to amplify samples using the following parameters: initial denaturation at 95°C for 10 minutes; 40 cycles of 95°C/10 seconds, 50°C/20 seconds, 75°C/15 seconds. Samples were then run through standard agarose gel electrophoresis to identify *B. burgdorferi* 16S rRNA amplification. Negative controls included uninfected zebrafish samples and no-RNA control for cDNA synthesis.

### Statistical analysis

Statistical analysis was performed using one-tailed paired Student’s t-test (Microsoft Excel, Redmond, WA, USA) on zebrafish survival rate and heart rates. Welch’s unpaired t-test (unequal sample sizes) was performed on 2 wpi zebrafish size (GraphPad Prism 8, San Diego, CA, USA). Statistical significance was determined based on p-values < 0.05.

## Results

### Zebrafish survival following immersion infection with *B. burgdorferi*

Following an 8-hour incubation period in either BSK-H media alone or a 1.0×10^7^ *B. burgdorferi*/mL suspension, zebrafish were removed from the suspensions, washed, and returned to the incubator at 28°C to recover for 72 hours in fish water. Survival rate was measured in both uninfected and infected zebrafish at 24, 48, and 72 hpi. For each of the time points, 100% survival was achieved for both uninfected and infected zebrafish (*n* = 120; Table 1). No statistical difference was observed between control and experimental groups, indicating that infection using 1×10^7^ spirochetes/mL of *B. burgdorferi* had no significant effect on zebrafish mortality.

**Table 1.** Zebrafish survival rate through 72 hpi for each variation of immersion infection. Each experimental condition used 20 zebrafish and was repeated between 3 and 6 times.

| Zebrafish<br>Developmental<br>Stage | *B. burgdorferi*<br>B31<br>concentration<br>(spirochetes/mL) | Number of<br>Repeats | Sample<br>Size, *n*<br>(total) | <u>Percent Survival (mean ± SD)</u> |  |  |
| --- | --- | --- | --- | --- | --- | --- |
|  |  |  |  | 24 hpi | 48 hpi | 72 hpi |
| 5 dpf | $1 \times 10^7$ | 6 | 20<br>(120) | $100 \pm 0\%$ | $100 \pm 0\%$ | $100 \pm 0\%$ |
| 5 dpf | $1 \times 10^8$ | 3 | 20<br>(60) | $33 \pm 10.4\%$ | $28 \pm 2.9\%$ | $25 \pm 5.0\%$ |
| 4 dpf | 1x10<sup>7</sup> | 3 | 20<br>(60) | 35 ± 8.7% | 30 ± 5.0% | 23 ± 2.9% |

We attempted a few variations to this above-mentioned immersion protocol to see if we can use more spirochetes for infection or use larvae less than 5-days-old. When the number of spirochetes was increased to 1×10^8^ spirochetes/mL in the immersion suspension, poor survival rates (33 ± 10.4% at 24 hpi; 28 ± 2.9% at 48 hpi; and 25 ± 5.0% at 72 hpi) were observed (*n* = 60; Table 1). Additionally, 4 dpf larvae were immersed with the concentration of 1×10^7^ spirochetes/mL, which also led to poor survival rate for each time point (35 ± 8.7%; 30 ± 5.0%; and 23 ± 2.9%, respectively; *n* = 60).

### Reduced heart rate in *B. burgdorferi*-infected zebrafish through 72 hpi

The next experiment was to measure heart rate in uninfected and infected zebrafish to see if *B. burgdorferi* infection had any detrimental effects on zebrafish stress levels. Zebrafish immersed with *B. burgdorferi* (Fig 2, blue bar) had significantly slower heart rates at 24 hpi as compared to zebrafish immersed in media only (p < 0.01). Heart rate of *B. burgdorferi*-infected zebrafish was 14.3% slower than that of uninfected zebrafish at 24 hpi. At 48 hpi, heart rate of infected zebrafish was 7.7% slower (p < 0.01) and at 72 hpi, infected zebrafish heart rate was 8.9% slower than uninfected zebrafish (p < 0.05).

**Figure 2.**
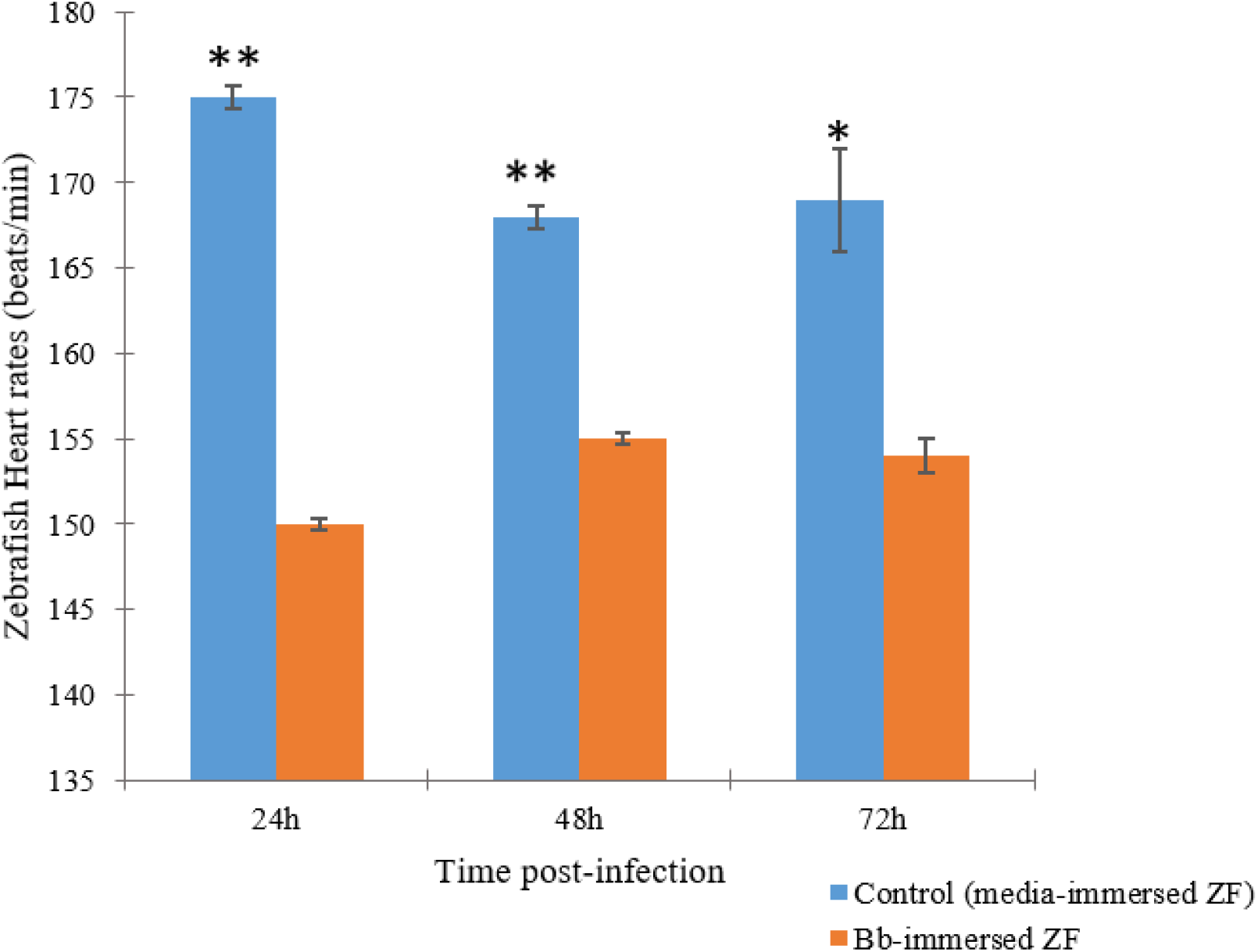
Heart rates of *B. burgdorferi*-infected zebrafish at 24, 48, and 72 hpi in immersion model. Error bars indicate the standard deviation of zebrafish heart rates from measurements performed across three repeats of the experiment (*n* = 15 for each time point); * p < 0.05; ** p < 0.01.

### *B. burgdorferi*-infected zebrafish are significantly smaller at 2 wpi

Throughout the 72-hour observation period post-infection, no morphological differences were seen between uninfected and infected zebrafish, and all fish showed alert, healthy behavior (data not shown). As an added measure, *B. burgdorferi*-infected zebrafish and uninfected control zebrafish were reared until 2 wpi followed by imaging to determine any differences in morphology, size, or behavior. Fig. 3 demonstrates a comparison of a control zebrafish (left) and a *B. burgdorferi*-infected zebrafish (right) maintained through 2 wpi. All zebrafish from this experiment (both uninfected and infected) were reared from the same clutch and were raised in exactly the same conditions. Measurements showed that infected fish were significantly smaller than control fish overall (p < 0.05). An average control fish measured 22.1 cm (± 2.7 cm) across the monitor, while infected zebrafish measured 15.2 cm (± 2.4 cm) (Fig. 4). Aside from control zebrafish being more biologically developed, both infection groups showed normal morphology (no dysmorphia) and healthy, alert behavior prior to anesthetization.

**Figure 3.**
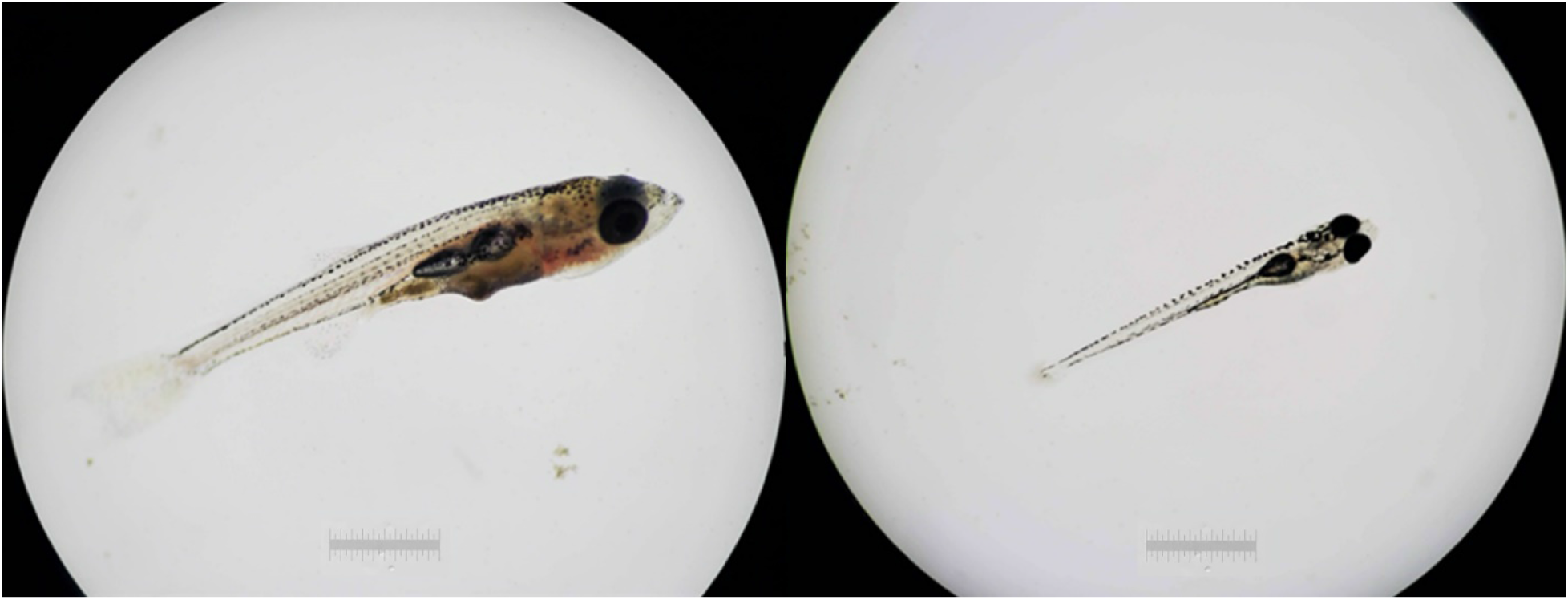
Representation of morphology and size of uninfected (left) versus *B. burgdorferi* infected zebrafish (right) at 2 wpi. Scale bar: 1 mm.

**Figure 4.**
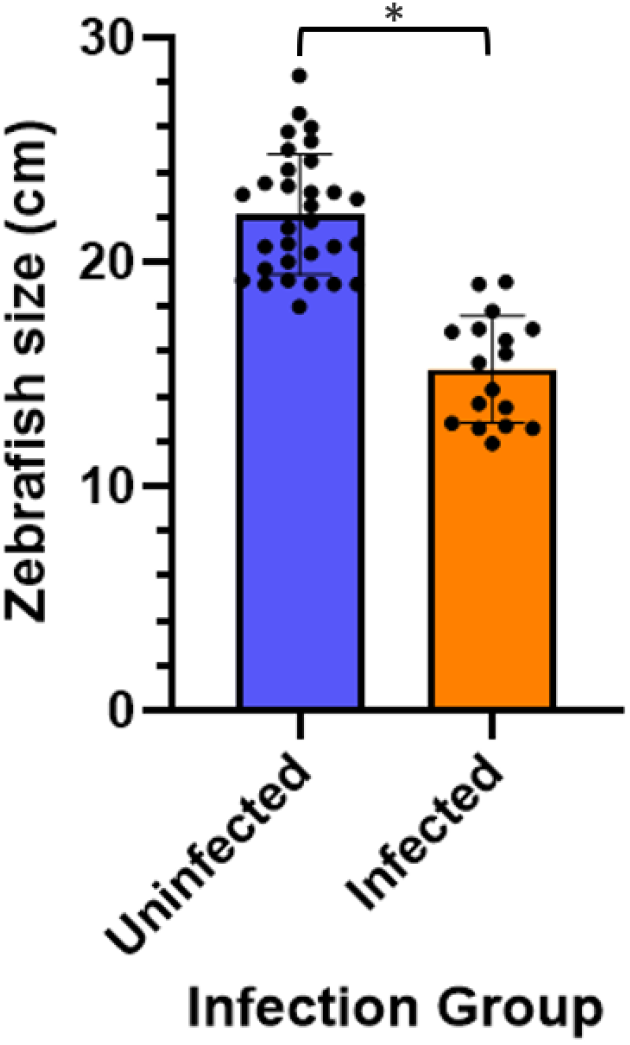
Size comparison of uninfected (blue) and infected (orange) zebrafish showing both mean ± 1 SD and individual fish sizes. Sample sizes *n* = 31 (uninfected), *n* = 17 (infected).

### *B. burgdorferi* spirochetes visualized in infected zebrafish via live fluorescence imaging

To ensure *B. burgdorferi* spirochetes were successfully introduced into the zebrafish system using the static immersion model, live fluorescence imaging was first performed at 24, 48, and 72 hpi on uninfected larvae and those infected with GFP-labeled *B. burgdorferi*. To ensure that there is no attached spirochete left on zebrafish and only spirochetes inside of fish can be detected, all fish are extensively washed as described in Material and Methods. Our results showed that spirochetes can be detected in the eyes (Fig. 5) and tail at 24 hpi. Since the suspension allowed spirochetes to be inhaled through the gills or ingested into the digestive tract, GFP aggregates were detected in the gills (Fig. 6, panel A) and stomach at 24 hpi. Of the *B. burgdorferi* GFP-immersed zebrafish imaged through 72 hpi (*n* = 60), 45% showed fluorescence in the eyes, gills, and tail (data not shown). PTU-treated, uninfected zebrafish which served as the negative control (*n* = 60), displayed no *B. burgdorferi* GFP fluorescence in any of their organs including the eyes, tail, or gills at any time points.

**Figure 5.**
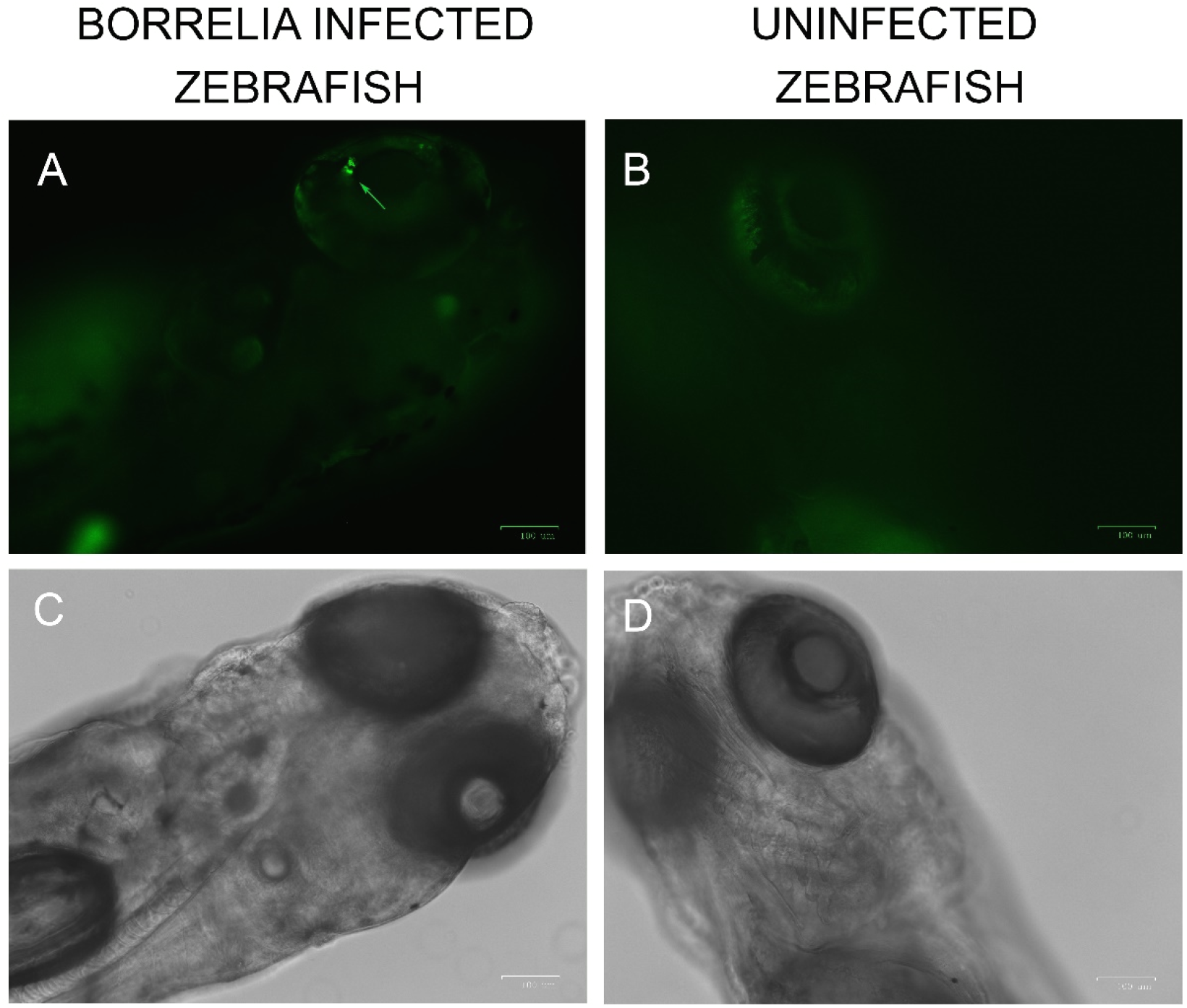
Comparison of *B. burgdorferi* GFP fluorescence visualized via live fluorescence microscopy between infected and uninfected zebrafish at 24 hpi. A) Spirochetes present in the eye of infected zebrafish under fluorescent filter. B) Uninfected zebrafish lacking fluorescence in eye. C, D) Corresponding brightfield images of infected and uninfected zebrafish, respectively. Scale bar: 100 μm.

**Figure 6.**
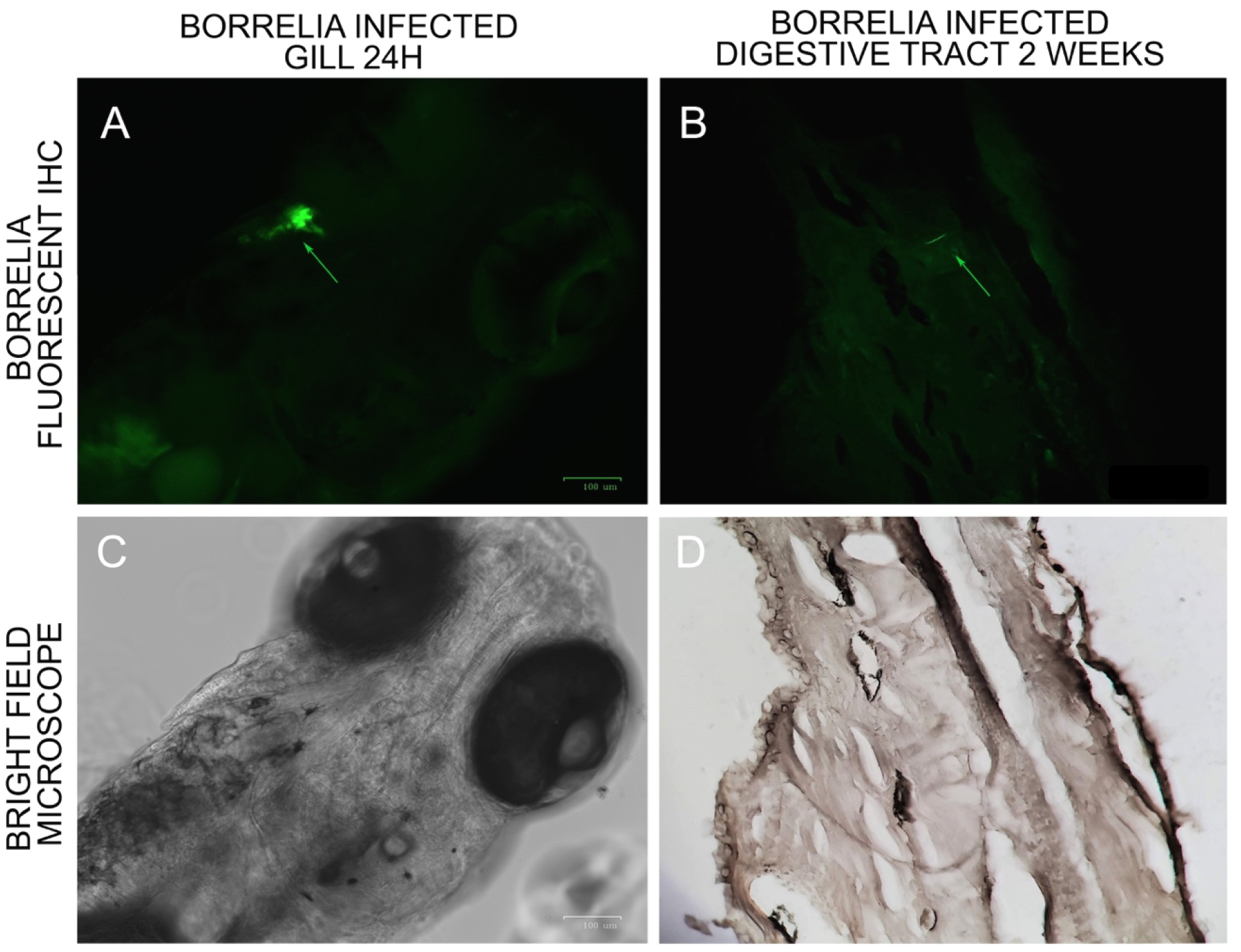
*B. burgdorferi* visualized in zebrafish gill and gut at 24 hpi and 2 weeks, respectively. A) Fluorescent image of *B. burgdorferi* GFP aggregate in zebrafish gill, C) Brightfield image of zebrafish larvae. Scalebar = 100 µm. B) Fluorescent image of *B. burgdorferi* spirochete in 2 wpi zebrafish digestive tract at 400x, D) Brightfield image of 2 wpi zebrafish.

Zebrafish which were reared to 2 wpi (uninfected and infected) were processed by paraffin embedding, sectioning, and staining with a *B. burgdorferi*-specific FITC labeled antibody followed by fluorescence imaging. Spirochetes were visualized in the zebrafish digestive tract (Fig. 6, panel B) and stomach. No fluorescence was visualized in any of negative control sections omitting *B. burgdorferi* FITC antibody, nor in uninfected control zebrafish (images not shown). Seven uninfected fish visualized across two experiments, none showed fluorescence, while 4 out of 6 of the infected zebrafish did stain positive *B. burgdorferi* (data not shown).

### Wholemount IHC reveals *B. burgdorferi* spirochetes in infected fish through 72 hpi

To further provide evidence that the immersion model can effectively infect zebrafish, wholemount IHC staining on infected 24 hpi, 48 hpi, 72 hpi zebrafish was utilized as described in Material and Methods (Fig. 7). Panels A, C, and E show IHC results using a *B. burgdorferi* monoclonal antibody (red stain), while Panels B, D, and F show brightfield microscopy images of the zebrafish. At different stages post-infection, *B. burgdorferi* spirochetes were found in different locations in zebrafish larvae. At 24 hpi, *B. burgdorferi* was found in the trunk muscle (Fig. 7, Panel A); at 48 hpi, *B. burgdorferi* was found in the overlying head epithelium and eye (Panel C); and at 72 hpi, *B. burgdorferi* was found in the heart and liver (Panel E), and tail. At 24 hpi, 83.3 ± 13.4% of infected zebrafish showed positive fluorescence; at 48 hpi, 100 ± 0 % showed fluorescence, and at 72 hpi, 83.3 ± 13.4% showed fluorescence. No fluorescence was seen in any of the control uninfected zebrafish for 24 through 72 hpi (*n* = 36; Table 2).

**Table 2.** Presence of *B. burgdorferi* in wholemount-stained uninfected and infected zebrafish.

| Immersion | *n*<sup>a</sup> | *n*<sup>b</sup> | % of zebrafish<sup>c</sup> (mean ± SD) |  |  |
| --- | --- | --- | --- | --- | --- |
|  |  |  | 24 hpi | 48 hpi | 72 hpi |
| *B. burgdorferi* | 3 | 4 | 83.33 ± 13.4% | 100 ± 0% | 83.33 ± 13.4% |
| BSK-H media | 3 | 4 | 0 ± 0% | 0 ± 0% | 0 ± 0% |

a Number of repeated experiments.
b Number of wholemount immunohistochemical stained zebrafish.
c Zebrafish visualized with *B. burgdorferi* infected.

**Figure 7.**
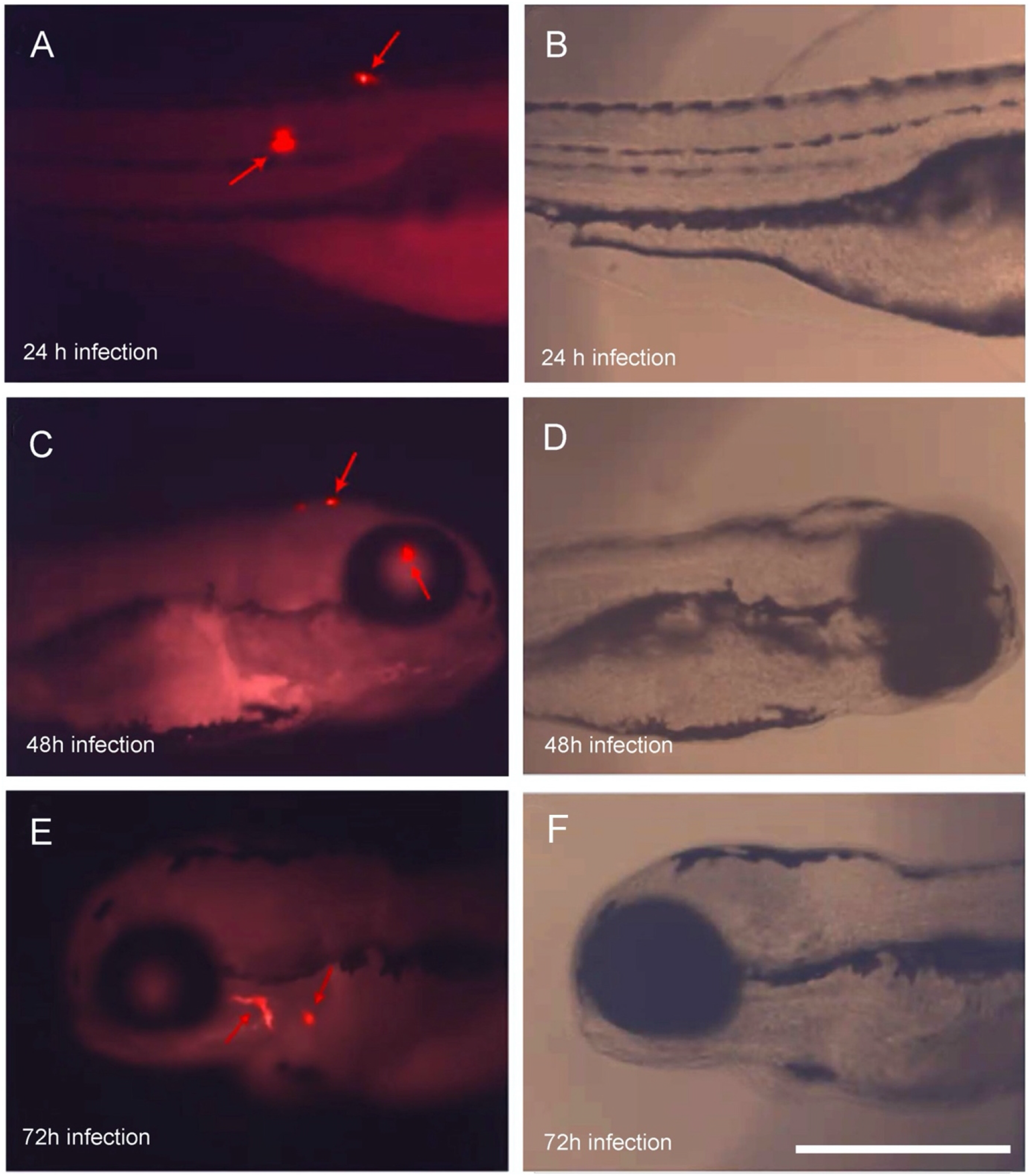
Wholemount IHC staining of *B. burgdorferi* 24 hpi, 48 hpi and 72 hpi zebrafish immersion model. A, C, E) Red fluorescent view of *B. burgdorferi*-infected zebrafish at 24, 48, and 72 hpi, respectively; B, D, F) Corresponding brightfield view of infected zebrafish at 24, 48, and 72 hpi, respectively. Scale bar = 250 µm.

### *B. burgdorferi* DNA detected through 72 hpi in infected zebrafish

Using a *B. burgdorferi* positive DNA control for comparison, it was demonstrated that all *B. burgdorferi*-infected zebrafish samples (four 24 hpi, three 48 hpi, and three 72 hpi) tested positive for *B. burgdorferi* DNA (Fig. 8) while none of the uninfected zebrafish and non template control samples showed positive amplification (*n* = 16). DNA detection for *B. burgdorferi* through 72 hpi was 92% (in total, *n* = 25 samples, each with 8 zebrafish larvae). A sample which had tested positive at 24 hpi was purified and sent for DNA sequencing and the obtained sequence was analyzed against the NCBI database. The sample returned a 99.9% identity match to *B. burgdorferi* senso stricto strain (data not shown).

**Figure 8.**
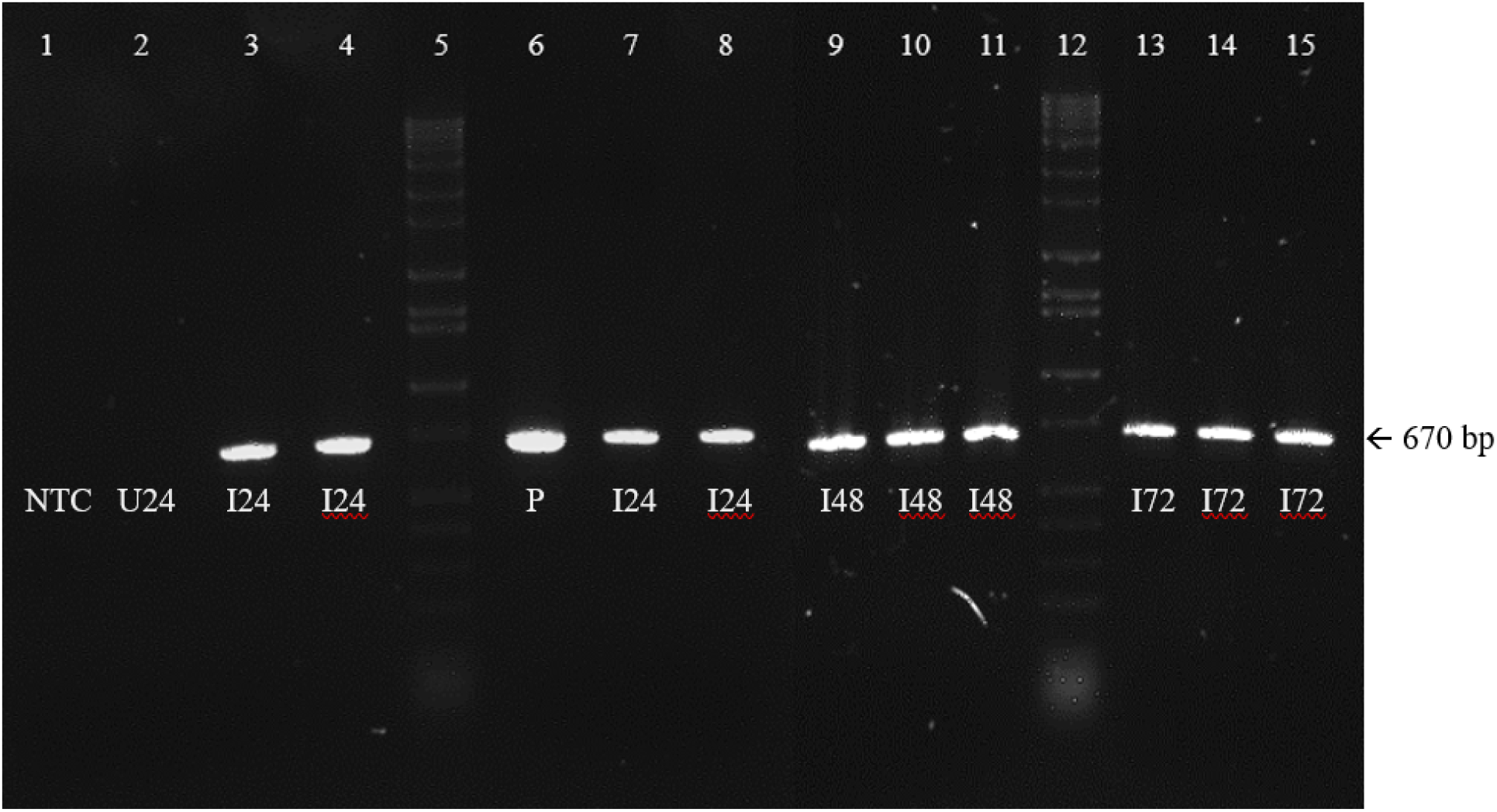
*B. burgdorferi*-amplified *PyrG* gene at 670 bp in *B. burgdorferi*-infected and uninfected zebrafish samples at 24, 48, and 72 hpi. Lane 1 shows the NTC; lane 2: 24-hour uninfected control (U24); lanes 5 and 12: Hi-Lo DNA ladder; lane 6: *B. burgdorferi* positive control (P); lanes 3-4, 7-8: 24-hour infected (I24); lanes 9-11: 48-hour infected (I48); lanes 13-15: 72-hour infected (I72).

### *B. burgdorferi* RNA detected through 10 days post-infection in infected zebrafish

Each day following immersion infection, RNA was extracted and cDNA was made from both uninfected and infected zebrafish and amplified for the *B. burgdorferi* 16S rRNA gene. Imaging of the obtained results revealed that all infected zebrafish samples from 1 dpi through 10 dpi were positive for *B. burgdorferi* (*n* =10), indicating replicating, live spirochetes (Fig. 9, lanes 4-13).

**Figure 9.**
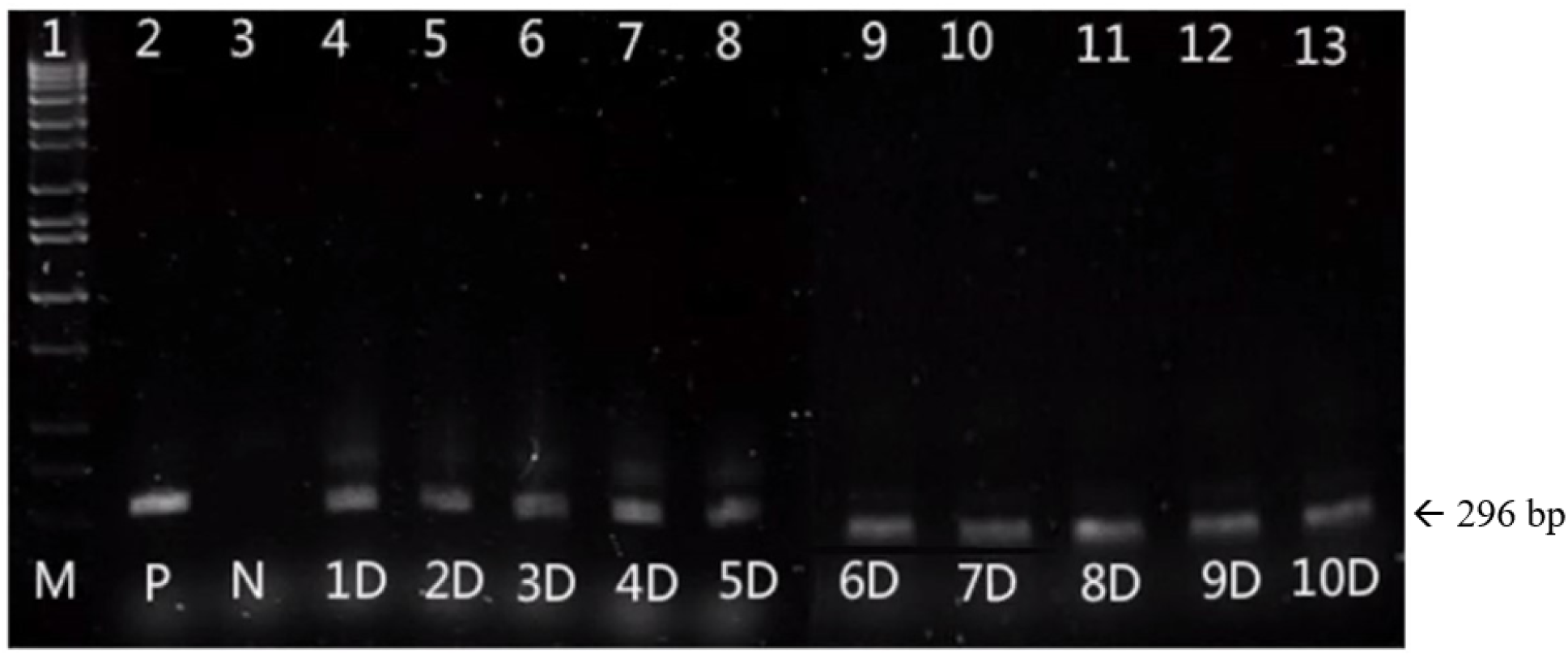
A representative gel electrophoresis image of *B. burgdorferi* 16S rRNA expression in the zebrafish immersion model at different times post-infection. Lane 1 shows Hi-Lo DNA ladder marker; lane 2: *B. burgdorferi* positive control (P); lane 3: uninfected zebrafish control (N); lanes 4-13: *B. burgdorferi*-infected zebrafish at 1 dpi through 10 dpi (1D-10D, respectively).

## Discussion

Given the lack of affordable, high-throughput models in Lyme disease research, this study aimed to investigate zebrafish (*Danio rerio*) as a novel *in vivo* model using a static immersion infection technique. The objectives were to develop straightforward, standardized infection procedures; maintain a high survival rate following infection; observe zebrafish for any morphological or behavioral changes post-infection; and detect *B. burgdorferi* inside zebrafish via PCR amplification and IHC visualization.

Our immersion infection technique has maintained survival and wellbeing of both zebrafish and *B. burgdorferi*. Survival of zebrafish through 72 hpi remained at 100% following infection in 1×10^7^ spirochetes/mL *B. burgdorferi* suspension (Table 1), indicating no toxicity from either the immersion suspension media (BSK-H) or from *B. burgdorferi*. Although suspension of zebrafish in pure water or PBS would tailor to the needs of the fish [34,35,36,37], *B. burgdorferi* have been found to form round bodies within 10 minutes of exposure to pure distilled water [38]. By diluting BSK-H media 1:1 with zebrafish water, the spirochete morphology of *B. burgdorferi* was preserved throughout the 8-hour infection period (verified via darkfield microscopy of the suspensions), indicating that *B. burgdorferi* remained in their active, motile state. Regarding zebrafish, no external morphological changes nor developmental abnormalities, such as heart edema (seen in larvae infected with *Porphyromonas gingivalis* as early as 24 hpi) or external lesions, were observed throughout the 72-hour observation period post-infection [39].

Though *B. burgdorferi*-infected zebrafish did not exhibit significant morphological changes, they did exhibit physiological changes through 72 hpi, shown by a significant decrease in heart rate compared to uninfected zebrafish. To our best knowledge, this response has not yet been reported in previous studies involving bacterial infection in zebrafish. Typical responses to infection might include erratic movement patterns [40], increased respiration rate due to bacterial burden [41], and an immediate-early immune response shown by increased inflammatory markers, such as those seen with *Edwardsiella tarda, Escherichia coli*, and *Listeria monocytogenes* infections [42,43,44]. Similarly, infection of zebrafish with the tick-borne pathogen, *Bartonella henselae*, showed increased expression levels of IL-1β, IL8, and VEGF [45]. However, these responses have not yet been studied in *B. burgdorferi-*infected zebrafish. In our model, we suspect the stress of a sustained *B. burgdorferi* infection induces detrimental effects on the zebrafish system, exhibited by the small, underdeveloped physique of infected larvae compared to control uninfected larvae at 2 wpi.

Visualizing *B. burgdorferi* is one major step to verifying the successful infection of zebrafish larvae. GFP-expressing bacteria have been utilized to study bacterial dissemination in zebrafish following immersion infection. For example, *Vibrio anguillarum* has been visualized in the mouth and gastrointestinal tract of zebrafish [36] and *L. monocytogenes* visualized in the gut lumen for a brief period after infection [44]. Our research has proven the presence of *B. burgdorferi* spirochetes inside the zebrafish following short-term infection through various techniques. First, zebrafish infected with GFP-Borrelia showed fluorescent spirochetes present in the eyes, gills, and tail through 72 hpi. Fluorescence signals were also visualized inside the stomach and digestive tract between 24 and 72 hpi (data not shown). Following the 8-hour immersion period, all zebrafish were washed extensively to wash away any exterior spirochetes, bringing us to the conclusion that those spirochetes visualized by fluorescence microscopy were indeed in the zebrafish tissue.

To further support the finding of spirochetes inside the zebrafish, *B. burgdorferi* were visualized by wholemount IHC staining of uninfected and infected zebrafish at 24, 48, and 72 hpi. Results supported what we found with live fluorescence imaging, where spirochetes were seen in the zebrafish eye, though also shown in the notochord, heart and liver. These additional locations could have been highlighted due to the orientation of the zebrafish when imaging, or the heart and liver being in close proximity to the zebrafish stomach where spirochetes could migrate through the soft tissue of the young larvae.

To ensure zebrafish were not able to clear *B. burgdorferi* from their system, uninfected and infected zebrafish were reared for 2 weeks post-infection then processed by paraffin embedding and FITC staining with a *B. burgdorferi*-specific fluorescent antibody. Not only do the numerous washes of the embedding process ensure any remaining spirochetes are removed from the outer surface of the larvae, but the slicing of tissue and visualization of any fluorescently stained spirochetes serve as additional proof of *B. burgdorferi* inside the zebrafish. At this timepoint, we found spirochetes in the stomach and digestive tract, indicating that spirochetes had not been destroyed 23through ingestions/digestion during the two weeks post-infection. This could suggest not only active, sustained infection, but also the potential for *B. burgdorferi* to migrate through the various zebrafish tissues for chronic, persistent infection, which is thought to begin around 2-3 wpi in mice [46].

Our final result supporting persistent *B. burgdorferi* infection was demonstrated by the presence of *B. burgdorferi* 16S rRNA harbored by zebrafish all the way through 10 dpi. The presence of fluorescent spirochetes visualized inside zebrafish at 72 hpi and detection of RNA activity through 10 dpi demonstrates the resistance of *B. burgdorferi* to host immune clearance, indicating the potential for long-term infection studies. Additionally, the detection of spirochetes on two separate levels (and no detection in uninfected zebrafish) shows successful infection of zebrafish larvae using the static immersion technique.

In conclusion, our results demonstrate that zebrafish serve as a promising animal host for *B. burgdorferi* infection which inspires future Lyme disease antimicrobial treatment studies. Compared to other models, zebrafish will increase the affordability of Lyme disease research, increase the number of sample sizes for improved significance, and provide more realistic results than *in vitro* models to provide a means of innovation in Lyme disease therapies.

## Acknowledgements

This work was supported by a grant from NASA Connecticut Space Consortium grant to EM. The authors thank for additional supports from National Philanthropic Trust, Joshua Foundation, Lyme Warriors, LivLyme Foundation for the research reported in this paper. The authors also thank University of New Haven, College of Art and Sciences for the establishment and maintenance of the Zebrafish Core Facility and Dr. Carter Takacs for his training to EM and MZ. The supporters had no role in study design, data collection and analysis, decision to publish, or preparation of the manuscript.

## Notes

### Competing Interest Statement

The authors have declared no competing interest.

